# Development of a new faecal DNA extraction method (HV-CTAB-PCI) for amplification of mitochondrial and nuclear markers used in genetic analyses of dugongs (*Dugong dugon*)

**DOI:** 10.1101/2022.11.24.517804

**Authors:** Vicky Ooi, Lee McMichael, Margaret E. Hunter, Aristide Takoukam Kamla, Janet M. Lanyon

## Abstract

Non-invasively collected faecal samples are an alternative source of DNA to tissues, that may be used in genetic studies of wildlife when direct sampling of animals is difficult. Although several faecal DNA extraction methods exist, their efficacy varies between species. Previous attempts to amplify mitochondrial DNA (mtDNA) markers from faeces of wild dugongs have met with limited success and nuclear markers (microsatellites) have been unsuccessful. This study aimed to develop a new tool for sampling both mtDNA and nuclear DNA (nDNA) from dugong faeces by modifying approaches used in studies of other large herbivores. First, amplification success of genetic markers from dugong faeces was compared between an established QIAamp and a newly developed DNA extraction method. Faecal DNA extracted using a new ‘High Volume-CTAB-PCI’ (HV-CTAB-PCI) method was found to achieve comparable amplification results to extraction of dugong skin DNA. As most prevailing practices advocate sampling from the outer surface of a stool to maximise capture of sloughed intestinal cells, this study compared amplification success of mtDNA between the outer and inner layers of faeces, but no difference in amplification was found. Assessment of the impacts of faecal age or degradation on extraction, however, demonstrated that fresher faeces with shorter duration of environmental (seawater) exposure amplified mtDNA and nDNA better than eroded scats. Using the HV-CTAB-PCI method, nDNA was successfully amplified for the first time from dugong faeces. This novel DNA extraction protocol offers a new tool that will facilitate genetic studies of dugongs and other large and cryptic marine herbivores in remote locations.

## Introduction

Genetic variation between individuals frequently influences the evolutionary resilience of species [1, 2]. Small populations lacking in gene flow have low genetic diversity, which may restrict their ability to adapt to environmental changes, leaving them vulnerable to extinction [3]. As threatened wildlife species dwindle in number, studies addressing genetic variation within and between populations become crucial for continued conservation efforts. To obtain critical genetic information on free-ranging wildlife, most studies rely on direct sampling of body tissues (e.g., blood, skin) as a source of high quality DNA [4]. However, there may be logistical and/or ethical challenges in direct sampling of large, rare, cryptic, or elusive species [5]. In contrast, non-invasive sampling through collection of animal traces, such as faeces, shed hairs, or sloughed skin as a source of DNA, mitigates the need to invasively capture, restrain or in some cases, directly observe the target wildlife [6]. One drawback of sampling biological traces is that quantity and quality of DNA may be compromised [6]. However, if collected and processed appropriately, non-invasive samples may provide comparable genetic information as obtained through direct sampling [e.g., 7].

Utilisation of faeces as an alternative source of DNA has its challenges. Apart from a generally low yield of amplifiable target DNA, faeces may contain PCR inhibitors that originate from digestive contents or the environment in which they are voided [8]. DNA from herbivore faeces is often more difficult to amplify compared to that of carnivores, since these may contain plant secondary metabolites that inhibit PCR [9]. The use of faecal DNA for genetic studies of marine species is even more restricted due to degradation and contamination in seawater [10]. Furthermore, the age of faecal samples may impact retrieval of target DNA, i.e., higher yield and quality of target DNA may be extracted from fresh compared to old scats [11].

The bulk of mammalian faeces comprises water and soluble molecules such as mucin, and the remainder consists of undigested food and microflora [12]. A small portion of faeces comprises epithelial cells sloughed from the host’s gastrointestinal tract [13]. Based on the assumption that sloughed cells would adhere to the surface of the egesta as it passes through the gastrointestinal tract, some sampling protocols swab the surface of scats to obtain these cells [14], whilst others resort to either homogenising the faecal sample or scraping, peeling, and/or washing its surface [8]. Surprisingly, the distribution of target cells and thus host DNA within faeces appears to have remained underexplored despite increasing efforts to isolate DNA from scats. Although various extraction methods have been used to retrieve host DNA from faecal samples, there is no ‘one-size-fits-all’ approach that works optimally for all species. Some widely used extraction techniques include the QIAamp Fast DNA Stool Mini Kit (hereafter referred to as QIAamp) [7, 15], phenol-chloroform-isoamyl alcohol (PCI) [11, 16] and cetyltrimethyl ammonium bromide (CTAB) [17]. Interestingly, the 2CTAB/PCI protocol has demonstrated success in extracting DNA from herbivore faeces [18].

Dugongs (*Dugong dugon*) are vulnerable marine mammals that are declining throughout their range from East Africa to Vanuatu [19]. Dugongs are challenging to study as they often reside in inshore, turbid waters and remote locations where access may be limited [20]. Dugongs spend most of their lives underwater, and their short surface intervals make them difficult to approach and sample directly [21]. Consequently, few genetic data are available for dugongs throughout much of their range. Of all regions supporting dugongs, northern Australia has been best studied [22] due to relative accessibility of some locations by researchers and intermittent recovery of carcasses. Conversely, in many regions outside Australia, dugong densities are low and variable; some populations have been extirpated, some are functionally extinct [23], and most are in decline [19]. These challenges to studying dugongs has rendered a non-invasive DNA sampling approach, i.e., via faeces, increasingly desirable as it may enable population genetic studies in areas where direct sampling is impracticable.

Early genetic studies of dugongs analysed mtDNA from recovered carcasses [21, 24]. Maternally-inherited mtDNA has high mutation rates, making it an effective marker for distinguishing deep taxonomic relationships and broad population structure [25]. Although mitochondrial studies suggested two dugong maternal lineages in Australia (i.e., north-western, and eastern populations), separated at Torres Strait [21], it was only when nDNA was used that finer scale population structuring was found within southern Queensland [26]. A more recent study evaluated population structure of dugongs along the entire eastern Queensland coast, using the same tandemly repeated microsatellites, and found an abrupt genetic break at the Whitsundays region, effectively separating northern and southern clusters [27]. Two less distinct subclusters were found within each of these main clusters, but a lack of tissue samples along the more remote northern coast limited discernment of population structuring along the entire coast. These patterns were also supported by analysis using 464 highly discriminatory single nucleotide polymorphisms (SNPs) [27]. A SNP represents variations of a single nucleotide base within the genome of different individuals, where the least abundant allele or nucleotide form occurs in more than 1% of the population [28]. Although a greater number of SNPs are required to provide the same resolution as microsatellites due to the biallelic nature of SNPs as opposed to the multiallelic nature of microsatellites, fewer genotyping errors associated with SNPs and their abundance throughout the genome have led to an increase in their use [29]. More importantly, the small target regions for SNP amplifications would allow for higher PCR amplification success in more degraded DNA such as those from faeces [30].

In areas where dugong tissues are difficult to obtain, faecal samples have been collected, although their use remains limited as only mtDNA has been successfully amplified, despite efforts to retrieve nDNA [21, 27]. Extraction of nDNA is more challenging because target DNA from faeces is often highly degraded, and the amount of available nDNA is less than that of mtDNA, since a diploid cell may contain many mitochondria but only possesses two copies of the nuclear genome [31]. However, a recent study has successfully extracted both nDNA and mtDNA from faeces of manatees, phylogenetically related to dugongs, using the QIAamp method with modifications including an additional purification step that removes PCR inhibitors [32]. Manatees are generalist herbivores that consume highly fibrous and abrasive macrophytes [33], whereas dugongs are seagrass specialists that preferentially graze on seagrasses with high nitrogen and low fibre content [34, 35]. As dugongs have high apparent digestibility of low fibre seagrass [36], they produce less fibrous faeces compared to manatees [32]. Although these sirenians possess some differences in terms of diet and digestive function [37], it is likely that the extraction methods used for manatees may work in dugongs since their less fibrous diets may reduce the occurrence of PCR inhibitors in their stools.

The major aim of this study was to develop and validate a faecal DNA extraction protocol and improve faecal sampling methodologies to enable successful amplification of nuclear and mitochondrial markers used in population genetic studies of dugongs. The main hypothesis was that mtDNA control region and nDNA, represented by a sex marker and ten SNP markers, could be amplified from the outer surface layers of a fresh dugong stool. The first objective was retrieval of sufficient target DNA from dugong faeces, using an optimal DNA extraction method, for consistent amplifications of zinc finger region (ZFX) within the X-chromosome of nDNA and the control region of mtDNA. The second objective investigated whether amplification success of both markers and the amount of target DNA differed between the outer versus inner layers of faeces with different levels of environmental exposure. The third objective sought amplification of ten previously developed SNPs [27], from fresh dugong faeces, using the optimal DNA extraction protocol. A representative SNP was also amplified in dugong faeces of different environmental exposure levels (EELs) to explore the impacts of degradation on SNP amplification success.

## Methods

### Field sampling

Two types of dugong faeces were collected: *Ex-ōceanum* faeces were those that were collected when dugongs were held out of water and were thus uncontaminated by seawater, whilst *in-ōceanum* faeces were those that were collected from the ocean and were contaminated by seawater.

Matched *ex-ōceanum* faeces (n= 8) and skin tissues (n= 8) were collected from live wild dugongs during the 2018 annual health assessment program in Moreton Bay, Queensland (27.4°S, 153.2°E), Australia [38]. Dorsal skin tissues were sampled using a skin scraper [39], and freshly voided *ex-ōceanum* faeces were collected onto a clean frisbee placed under the anus of each dugong held on deck [38]. Frozen *ex-ōceanum* faecal samples collected in 2016 (n= 8) and 2017 (n= 8) were also used in this study. *Ex-ōceanum* faeces represent the freshest and least degraded of all faecal types and were categorised under Environmental Exposure Level 1 (*ex-*EE-L1). The surfaces of these faeces were textured, ranging from yellow-brown, or light brown to dark brown, and the inner core was usually more dough-like as opposed to fibrous with yellowish pigmentation.

*In-ōceanum* faeces of four different EELs or exposure states were collected from the ocean at different time points post-elimination. Fresh *in-ōceanum* faeces with exposure level 2 (*in-*EE-L2) (n= 8) were collected from the benthos through free-diving immediately as the feeding herd of dugongs was seen leaving an area (estimated at < 1 h post-elimination).

Floating faeces of exposure level 3 (*in-*EE-L3) (n= 8) were collected from the ocean surface in an area where the feeding herd was spotted nearby (∼ 2 – 4 h post-elimination). The *in*-EE-L2 and *in*-EE-L3 faeces have comparable morphologies: cylindrical (large calibre) of variable length. Their inner core usually presents comparable to *ex*-EE-L1; however, fibre proportion can be variable. Their colours are usually dark to light brown. Exposure level 4 (*in-*EE-L4) (n= 8) faeces were collected from the ocean surface where individuals were spotted foraging at a distance (∼ 5 – 7 h post-elimination). Morphologically, they are usually smaller and/or shorter than *in*-EE-L2 and *in*-EE-L3, though large specimens have been found. The shape of *in*-EE-L4 faeces remains cylindrical, yet jagged edges and lumps emerge in this category. Their colours are darker brown to black on the surface and the inner core are more yellowed with a slightly more fibrous nature. The most eroded faeces, exposure level 5 (*in-* EE-L5) (n= 8), were collected from the ocean surface in an area where no dugongs were spotted (possibly > 7 h post-elimination). The *in*-EE-L5 faeces tend to be in small pieces as they are broken down by ocean currents and/or coprophagous animals since they were exposed to the environment for the longest duration. The eroded faeces commonly appear charred black or dark brown in colour, and the inner core is mostly highly fibrous, though in some cases, they retained less fibre. As they are found in small pieces, the shape can differ from being cylindrical to round-like. (Summary: Table 1, Description: Fig 1).

**Fig 1.**
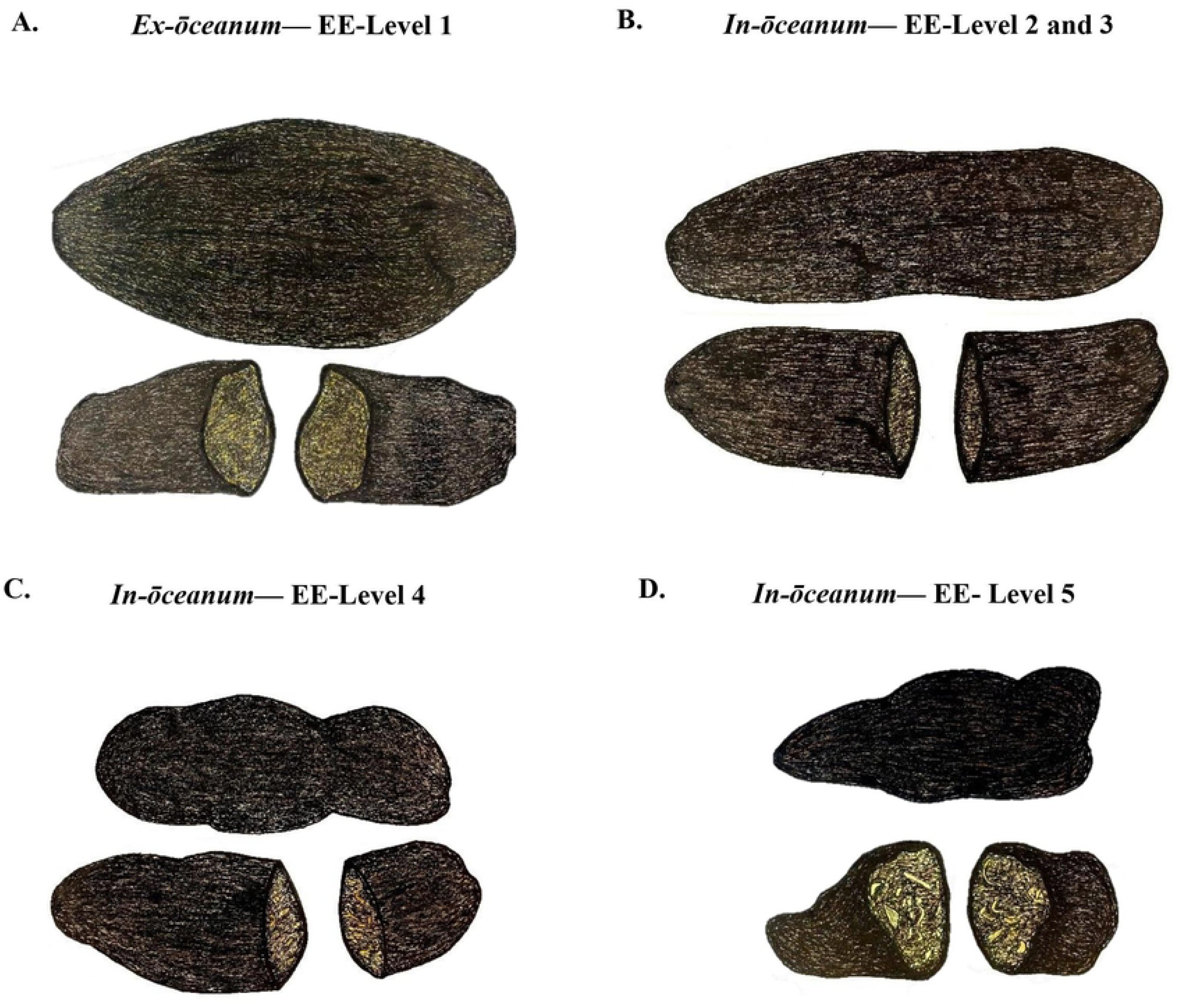
Classification of dugong faeces into five categories as illustrated with an above view (top) and transverse cross-section (bottom). Comparing freshly excreted samples on deck to those most aged in the ocean, a respective transition of surface colour was observed from lighter to darker, yellow-brown, or light brown to black, while sizes trended from large to small, and core composition generally retaining higher fibre proportion over time. (A) *Ex-ōceanum* faeces (*ex*-EE-L1), originating beneath the dugong, were flattened in a shallow frisbee. Thus, the above view displays a wider shape (top) and a flatter profile from the side (bottom). (B) *In-ōceanum* Level 2 and 3 faeces. (C) *In-ōceanum* Level 4 faeces. (D) *In-ōceanum* Level 5 faeces.

**Table 1.**
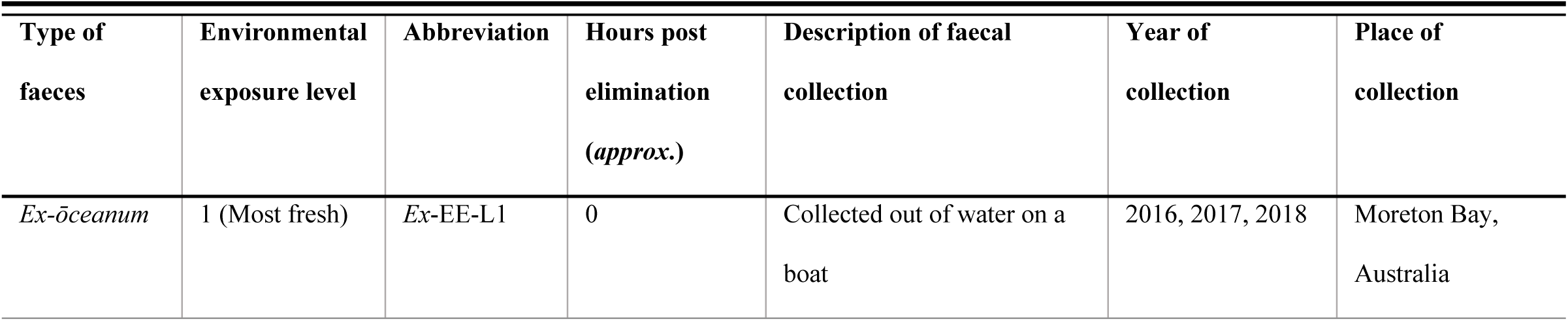
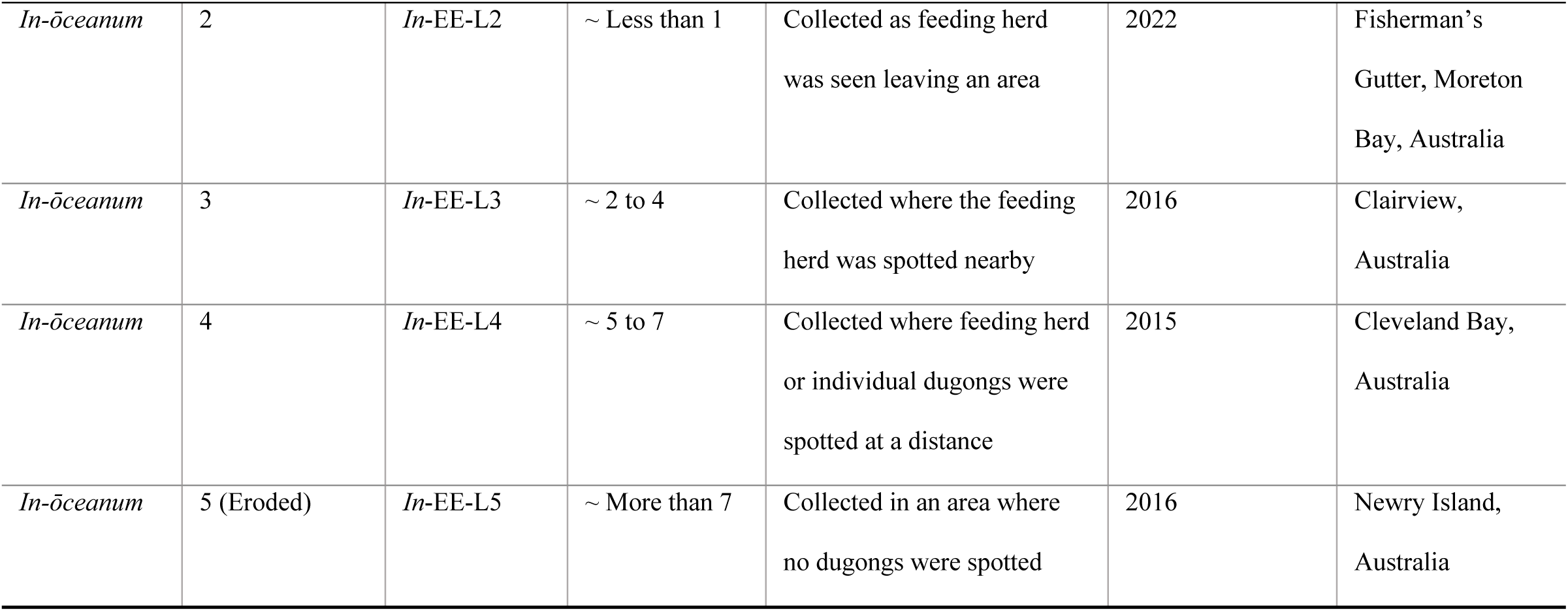
Summary on the description and categories of all faecal samples collected and used in this study, including the year and place of collection.

Faeces were placed in a ziplock bag upon retrieval, stored in an esky in the field, and later frozen at -20 °C. Skin tissues and a single liver tissue sample from a neonate dugong, were used in this study as positive controls; both mtDNA and nDNA have been previously extracted from these tissue types [26].

### Experimental Design

To develop an appropriate protocol for DNA extraction from dugong faeces, two faecal sampling and two processing techniques were compared, and the approach that recovered the highest amount of total (dugong and exogenous) DNA and target (dugong) mtDNA was used in all further extractions. Two DNA extraction methods, i.e., the QIAamp method and a newly developed ‘High volume-CTAB-PCI’ (HV-CTAB-PCI) method, were tested. Total DNA concentration extracted using each protocol, and the amplification success (including each of PCR and triplicate success) of mtDNA control region and ZFX (sex marker) were compared (S1 Fig A). PCR success represents the percentage of samples amplified, while triplicate success represents the number of replicates amplified out of three technical replicates for each sample.

The QIAamp method was used to extract DNA from the outer surface and inner core of dugong faeces of four different EELs (all except *in*-EE-L2 faeces). The amount of DNA used in qPCR reactions was normalised to enable comparisons between faecal samples. Total DNA concentration, relative amount of dugong mtDNA extracted, and amplification success were compared under a two-factorial design (S1 Fig B). The HV-CTAB-PCI method was used to extract DNA from dugong faeces of five different EELs, and both mtDNA control region and ZFX (nDNA) were amplified. Total extracted DNA concentration, and amplification success of each marker were compared between faeces of different EELs (S1 Fig B).

DNA extracted using the HV-CTAB-PCI method from *in*-EE-L2 faeces was used to trial the amplification of ten SNP primer sets, and amplification success was compared. One SNP marker was selected to be amplified using DNA extracted with the HV-CTAB-PCI method from all *in-ōceanum* faeces, and amplification successes were compared (S1 Fig C).

### Faecal sampling and processing

Using a sterile surgical blade, 220 mg of faecal material was scraped from the outer surface of 16 *ex-ōceanum* faecal samples, each weighed and stored in a 2 mL microcentrifuge tube (Note: For inner core sampling, the stool was broken in half and faecal material from the inner core of faeces was sampled). Each faecal sample was then either transferred into a sterilised mortar and ground into a fine powder with a pestle using liquid nitrogen (N_2_) (referred to as ‘scrape-and-grind’ technique) (n= 8) or left unground (referred to as ‘scrape-only’ technique) (n= 8) (Fig 2).

**Fig 2.**
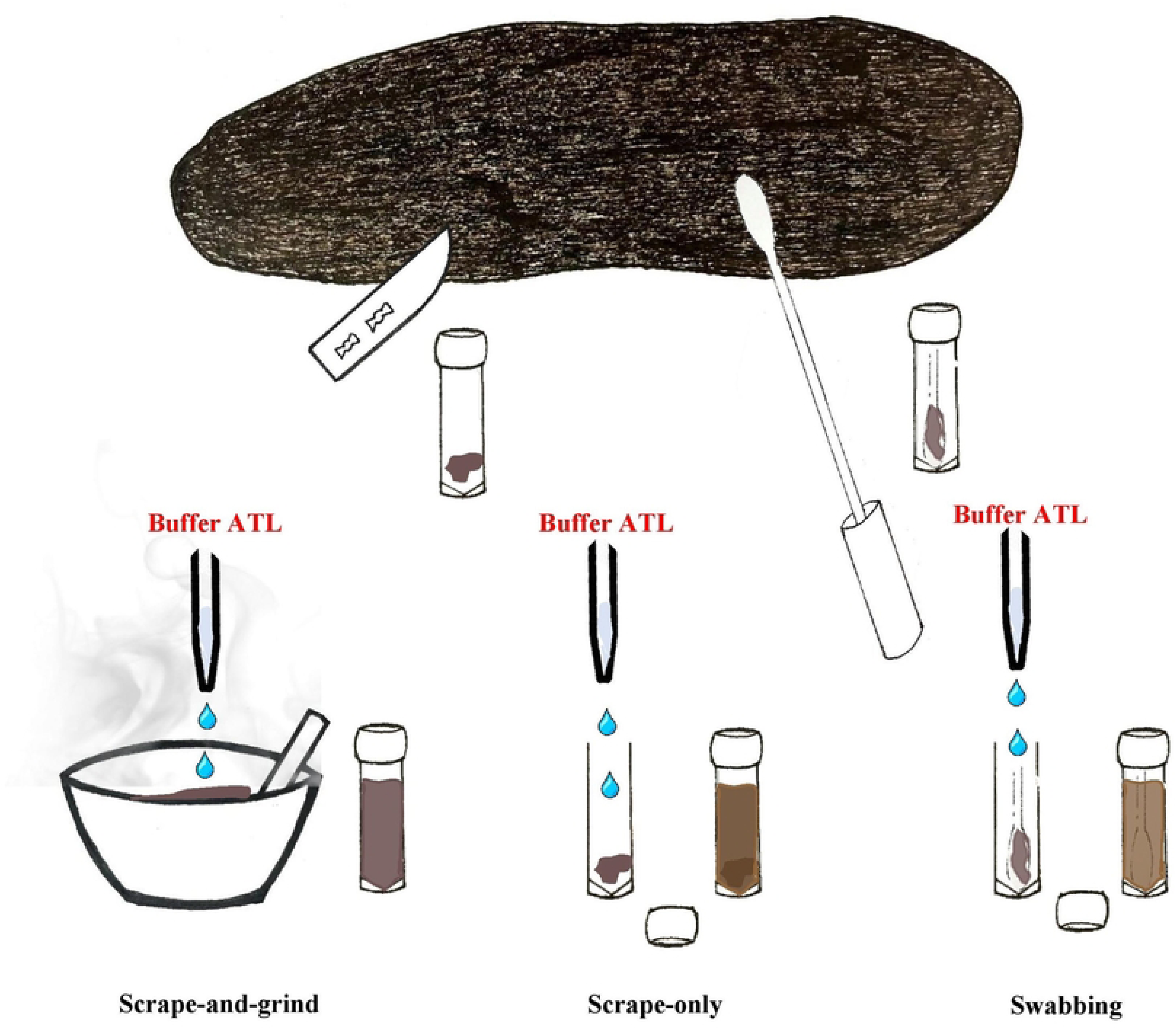
Faecal sampling and processing techniques used in this study. Faecal material was either scraped from the outer surface of faeces or swabbed along the entire faeces for DNA extraction. When faecal material was obtained via scraping, it was either further processed by grinding in liquid nitrogen prior to the addition of lysis buffer (Buffer ATL) or added directly with lysis buffer without any processing done. When faecal material was sampled using the swabbing technique, no further processing was performed.

A further eight *ex-ōceanum* faeces were sampled through swabbing the entire outer surface using a rayon swab (Copan, CA, USA) (referred to as ‘swabbing’ technique). Once the swab tip was entirely covered in faecal material, it was transferred into a 2 mL tube, and the swab’s shaft was trimmed to fit. No further processing was conducted (Fig 2).

### DNA extraction from faeces

#### QIAamp protocol

DNA was extracted using QIAamp Fast DNA Stool Mini Kit (#51604, Qiagen, Germany). When swabbing or scrape-only techniques were used, 1 mL of InhibitEX buffer was added into the 2 mL tube containing the faecal sample [32]. When scrape- and-grind technique was used, 500 μL of InhibitEX buffer was added to the mortar with ground faeces for further grinding prior to the transfer of faecal mixture back into the 2 mL tube, and another 500 μL was used to rinse the remaining faeces from the mortar into the same tube. The resultant mixture was vortexed for 1 min (Note: swab was removed after vortexing) and centrifuged for 1 min at 20,000 *g* (14,000 rpm) to pellet the stool particles. The remainder of the extraction was performed following manufacturer’s instructions, but by using 800 μL of supernatant from the lysate, and subsequently equal amount of Buffer AL and 95 % ethanol, with a final 100 μL elution volume.

#### HV-CTAB-PCI protocol

An extraction protocol was developed using the lysis buffers and general concept of the 2CTAB/PCI method [18]. Approximately 1 g of faecal material was processed using the scrape-and-grind technique, and 1 mL of Lysis Buffer 1 (LB1: CTAB 2 %, Tris– HCL 100 mM, EDTA 20 mM, NaCl 1.4 M, pH 7.5) was added to the mortar to further grind the powdered faeces before the mixture was transferred into a 15 mL tube. This was repeated twice to ensure that any remaining faecal material in the mortar was collected. Another 2 mL of LB1 was added into the same tube, making a total 5 mL LB1 added to the faecal sample. The mixture was vortexed and incubated at 60 °C for 3 h, with occasional mixing for cell lysis. After centrifuging at 3,150 *g* (4,000 rpm) for 12 min, 4 mL of supernatant was transferred into a new 15 mL tube and equal volume of phenol: chloroform: isoamyl alcohol (PCI, 21:20:1) was added to the supernatant, and gently mixed. The mixture was centrifuged as above, and 3 mL of the aqueous phase was transferred into a new 15 mL tube. Next, 330 μL of Lysis Buffer 2 (LB2: CTAB 10 %, NaCl 0.5 M, pH 5.5) was added to the supernatant, and incubated at 60 °C for 4 h, with occasional mixing, for further lysis. Thereafter, 104 uL of protease (#P5147, Sigma Aldritch, USA) was added to the lysate to digest proteins for 1 h at 60 °C. Then, equal volume (3434 μL) of PCI was added to the mixture, gently mixed, and centrifuged as above. Three mL of the aqueous phase was transferred into a new 15 mL tube and equal volume of isopropanol was added for overnight DNA precipitation at -20 °C. The sample was centrifuged for 20 min at 8000 *g* (5,200 rpm), and all supernatant was decanted. The pellet was washed once with 400 μL of 70 % ethanol. After being vortexed and centrifuged at 3,150 *g* for 12 min, the supernatant was decanted. The pellet was air dried at room temperature for 15 min and resuspended in 250 μL of TE buffer (10 mM Tris– HCl, 1 mM EDTA, pH 8).

### DNA extraction from skin

DNA was extracted using DNeasy Blood and Tissue kit (#69504, Qiagen, Germany). Approximately 10 mg of dugong skin tissue was ground to fine powder using liquid N_2_, and 180 μL of Buffer ATL was added into the mortar for further grinding. The mixture was transferred into a 1.5 mL tube, and the addition of another 50 μL of Buffer ATL to the mortar aided recovery of any remaining tissue. The rest of the extraction followed manufacturer’s protocol, using a 3 h incubation period, with 100 μL elution in a 1.5 mL tube. A centrifuging step was added following the addition of Buffer AL to remove precipitate, and ethanol was added to the supernatant. Using 2 μL of DNA extract, concentration (ng/μL) and purity (A260/280 and A260/230) of all extracted DNA were measured using NanoDrop™ 1000 Spectrophotometer (Thermo Fisher Scientific, USA). DNA isolates were stored at -80 °C until use.

### Real-time PCR assays

Quantitative polymerase chain reaction (qPCR) was conducted in a CFX96 Touch Real-Time PCR Detection System (Bio-Rad, USA), using 96-well PCR plates, and a ‘no template control’ was used to detect contamination. DNA extracted from tissue was used as positive control. Melt curve analysis was performed to determine melt temperature of the primers and to detect presence of non-specific amplifications.

#### mtDNA and ZFX markers

The control region of mtDNA and the ZFX region of nDNA were amplified using specific primers developed by Tol *et al* [40] and McHale *et al* [41], respectively (Table 2). These markers were used throughout the entire study except for SNP amplifications.

**Table 2.**
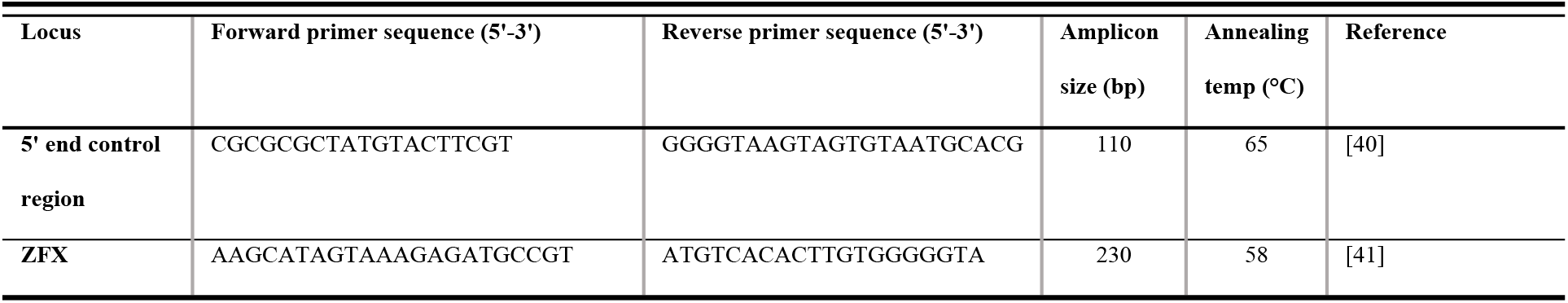
Forward and reverse primer sequences for mtDNA control region and ZFX markers used in this study, with their corresponding amplicon sizes and annealing temperatures.

PCRs were performed with a 20 μL reaction volume, each consisting of a variable amount of total DNA (Table 3), variable concentration of forward and reverse primers (Table 3), nuclease-free water, and 10 μL of PowerUp™ SYBR™ Green Master Mix (Applied Biosystems, USA).

**Table 3.** Summary of the total DNA (ng) and concentrations of forward and reverse primers (μM) added to the PCR reaction for the corresponding experiments and DNA extraction methods.

| DNA extraction method | Usage | Primer | Total DNA added (ng) | Concentration of forward and reverse primers used ($\mu\text{M}$) |
| --- | --- | --- | --- | --- |
| **QIAamp** | Compare faecal sampling techniques | mtDNA | 50 | 0.1 |
| **QIAamp** | Compare DNA extraction method<br>Compare faecal layer sampling on faeces of different quality | mtDNA | 100 | 0.1 |
| **QIAamp** | Compare DNA extraction method | ZFX | 100 | 0.5 |
| **HV-CTAB-PCI** | Compare DNA extraction method,<br>Compare faecal quality | mtDNA | Neat DNA extracted/2 | 0.1 |
| **HV-CTAB-PCI** | Compare DNA extraction method,<br>Compare faecal quality | ZFX | Neat DNA extracted/2 | 0.3 |
| **HV-CTAB-PCI** | SNP amplification | SNP | Neat DNA extracted/2 | 0.5 |

When DNA extracted with QIAamp method was used, the amount of total DNA added to the reactions was standardised and served as a normaliser to enable comparisons between samples. For the comparison of sampling from different layers of faeces of different EELs, a pilot experiment was conducted to determine whether 10 or 100 ng of total DNA should be used in the reactions (S2 Fig). One-hundred ng of total DNA was chosen because eroded faeces appeared to have higher PCR success using this amount, although the difference was not statistically significant. When DNA extracted with HV-CTAB-PCI method was used, a 1:2 dilution factor provided the most consistent amplification of all markers in the dilution experiments and was thus used for all subsequent PCR reactions.

Cycling conditions were 50 °C for 2 min, then 95 °C for 2 min, followed by 45 cycles of 95 °C for 15 s, respective annealing temperature (Table 2) for 1 min, and ending with a melt profile from 65 °C to 95 °C, with 0.1 °C increments.

#### SNP markers

The 10 SNP primers used were developed by McGowan [27] (Table 4). PCRs were performed as described previously. Total DNA diluted 1:2 was added to each reaction, and 0.5 μM of forward and reverse primers were used (Table 3).

**Table 4.**
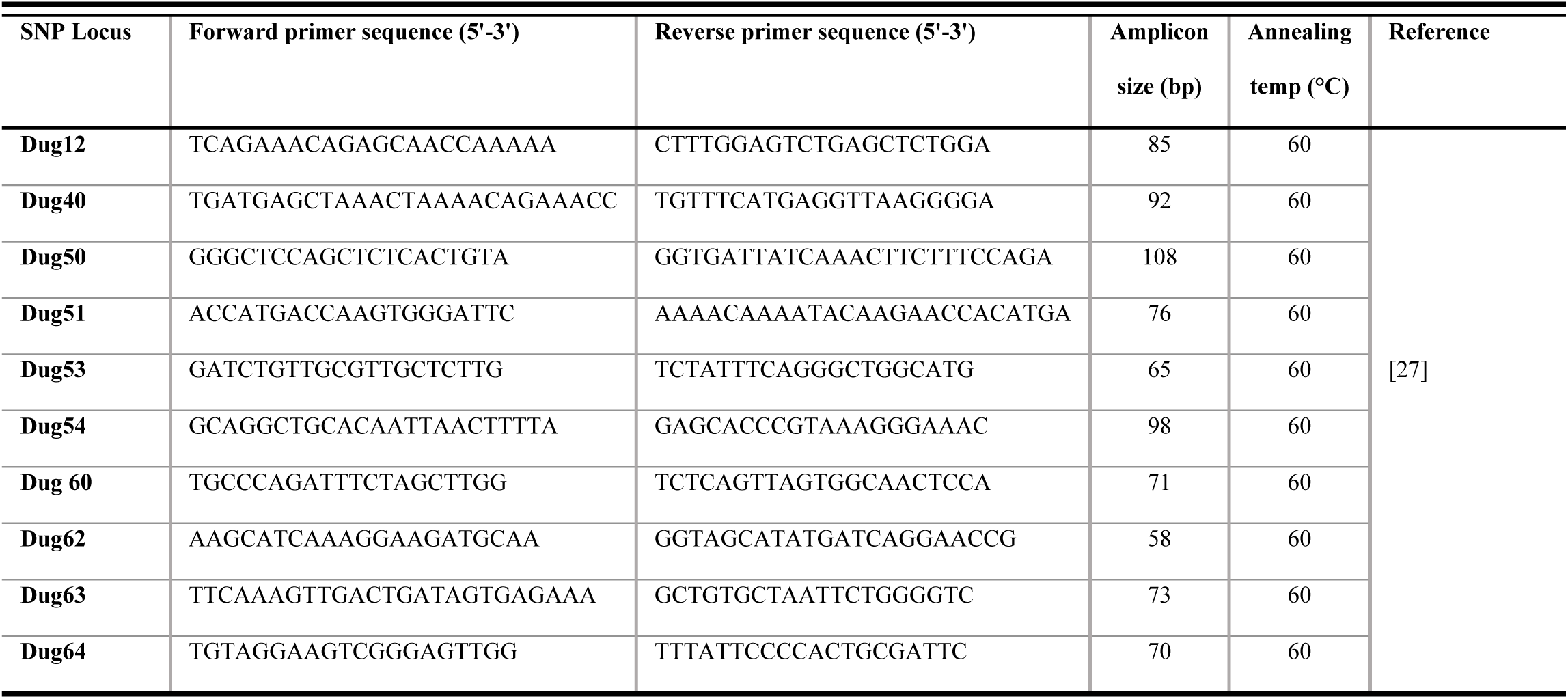
Forward and reverse sequences of SNP primers used in this study, with their corresponding amplicon sizes and annealing temperatures.

Cycling conditions were 50 °C for 2 min, 95 °C for 2 min, followed by 45 cycles of 95°C for 15 s, 60 °C for 30 s, 72 °C for 30 s, and 72 °C for 15 min. A melt profile from 65°C to 95 °C, with 0.5 °C increments was performed.

### qPCR intra-assay and inter-assay variability

PCR reactions were performed in triplicate to account for intra-assay variability, and the results were averaged for final analysis. To account for inter-assay variability, threshold cycle (C_T_) was manually set to the same value using the CFX Maestro software (Bio-Rad, USA) when C_T_ values were compared, and tissue control was used as an inter-plate calibrator in each plate when relative quantity was calculated and compared.

### Primer efficiency and linearity

A standard curve was created for both mtDNA and ZFX primers through dilution experiments, starting from 1000 ng of total DNA, with seven 10-fold dilutions for two skin DNA isolates. The linear dynamic range (LDR) and limits of detection (LOD) for each primer sets were determined from dilution curves, and PCR efficiency (*E*) was calculated from the slope of dilution curve as per Ruijter *et al* [42].

Five 10-fold serial dilutions and four 10-fold serial dilutions of two faecal DNA extracts, starting from 5 × neat DNA extracted were performed for mtDNA and ZFX primers, respectively. This was used to determine the most appropriate quantity of DNA extracted via the HV-CTAB-PCI method to be added to the PCR reactions.

### Primer specificity

Unpurified PCR products were submitted to the Australian Genome Research Facility (AGRF) at The University of Queensland, Australia, for dual-direction Sanger sequencing. Sequences from mtDNA and ZFX amplifications underwent BLAST searches on GenBank, and sequence similarity was used to confirm specific amplification of respective dugong DNA. The resultant SNP sequences were compared to amplicon sequences of McCarthy [43] and McGowan [27] to confirm amplification of interested regions, and to determine the SNP allele possessed by the individuals. Specificity of PCR amplicons was additionally verified through a melt curve analysis for each PCR reaction. SNP sequences were aligned and trimmed using MEGA11 software [44].

### Data processing

As C_T_ value is inverse to the amount of target DNA present within a sample, the inverse of C_T_ value was used for comparative analyses to facilitate visualisation of results. Relative quantity of target mtDNA was calculated in the CFX-Maestro software (Bio-Rad, USA), using a tissue sample as control and *E*. The relative quantity was multiplied by 100,000 for easier interpretation of small numbers. For samples that failed to amplify, a C_T_ value of the number of PCR cycles used + 1 was assigned to enable the determination of lowest possible C_T_ value difference for comparison.

### Statistical analyses

All statistical analyses were performed using *R* (version 4.2.0, [45]). For all parametric models fitted, diagnostic plots were made to check whether the assumptions for homogeneity and normality of residual variance were met. Normality was further confirmed with the Shapiro-Wilk’s test. Data were log-transformed when residuals were not normal. When significant difference was found in the initial model, a post-hoc pairwise Tukey test was performed using *lsmeans* package for further pairwise analyses. False discovery rate adjusted p-values were obtained when non-parametric tests were used. Graphs were created using *ggplot2* [46], *dplyr*, *hrbrthemes* [47], *Rmisc* [48], *ggpmisc* [49], *reshape*, *ggsignif* [50], and *ggpubr* [51] packages.

A linear mixed effects model (LMEM) from *nlme* package [52] was used to compare total DNA concentration and inverse C_T_ between different faecal sampling techniques, where random effects of individuals within PCR plates were accounted for. It was also used to compare log total DNA concentration recovered between different DNA extraction methods, accounting for the random effects of individuals between PCR plates. For all primers, PCR success (binary outcome: successful—if one or more amplified versus unsuccessful—none amplified) between DNA extraction methods was compared using Pearson’s Chi-square test (χ^2^), with a Monte Carlo simulation utilising 10,000 replicates, followed by post-hoc pairwise Chi-square tests. The number of amplifications within a triplicate (i.e., Triplicate success: 0, 1, 2, or 3) for all primers, was compared between DNA extraction methods using Kruskal-Wallis test and post-hoc pairwise Wilcoxon rank sum (*W*) test.

To determine the effects of environmental exposure and layers of faeces sampled on the amplification of dugong DNA, a full interactive generalised linear model (GLM) with all variables (sampling layer, faecal EELs, year of collection, individuals) was fitted to compare the total DNA concentration and relative quantity of dugong mtDNA. The Akaike Information Criterion (AIC) or stepAIC function from *MASS* package [53] was used to determine significant variables through a dual-direction stepwise model selection. A final additive LMEM was fitted to compare total DNA concentration and log relative quantity of dugong mtDNA between faecal sampling layers and faecal EELs, accounting for random effects of different individuals. The relationship between PCR success and log total DNA was tested using a quadratic GLM with binomial responses.

A likelihood ratio Chi-square test (LR-Chi-square test) was used to compare between interactive (sampling layer*faecal EELs) and additive (sampling layer + faecal EELs) GLMs, and a final additive GLM with biased reduction was fitted, using *brglm2* package [54], to compare PCR success between the two explanatory variables. A LR-Chi-square test was also used to compare between four multinomial log-linear models (from *nnet* package; [53]) fitted with interactive, additive, and single explanatory effects on the triplicate success of samples. A post-hoc pairwise *W* test was then used to compare triplicate success between sampling layer and faecal EELs.

When HV-CTAB-PCI method was used, total DNA concentration was compared between faecal EELs using a one-way analysis of variance (ANOVA), whilst PCR and triplicate successes were compared between different faecal EELs using the χ^2^ and Kruskal-Wallis tests as described earlier. Similarly, two non-parametric tests were also used to compare the PCR and triplicate successes between different primers and faecal types in SNP amplifications.

### Ethics Statement

Dugong samples were obtained under The University of Queensland Animal Ethics Permit SBS/181/18, Scientific Purposes Permit WISP14654414, Moreton Bay Marine Parks Permit MPP18-001119, and Great Barrier Reef Marine Park Permit G14/36987.1.

## Results

### PCR efficiencies, LDR and LOD

PCR efficiency of tissue DNA was 96.3 % and 86.2 % for mtDNA and ZFX primers, respectively (Fig 3A). All amplifications were within LDR (1-1000ng of total DNA) and LOD (0.001 ng of total DNA) for both primers. The *E* of faecal DNA indicated slight inhibition (mtDNA= 106.3 %, ZFX= 104.7 %), which was addressed through dilutions (Fig 3B).

**Fig 3.**
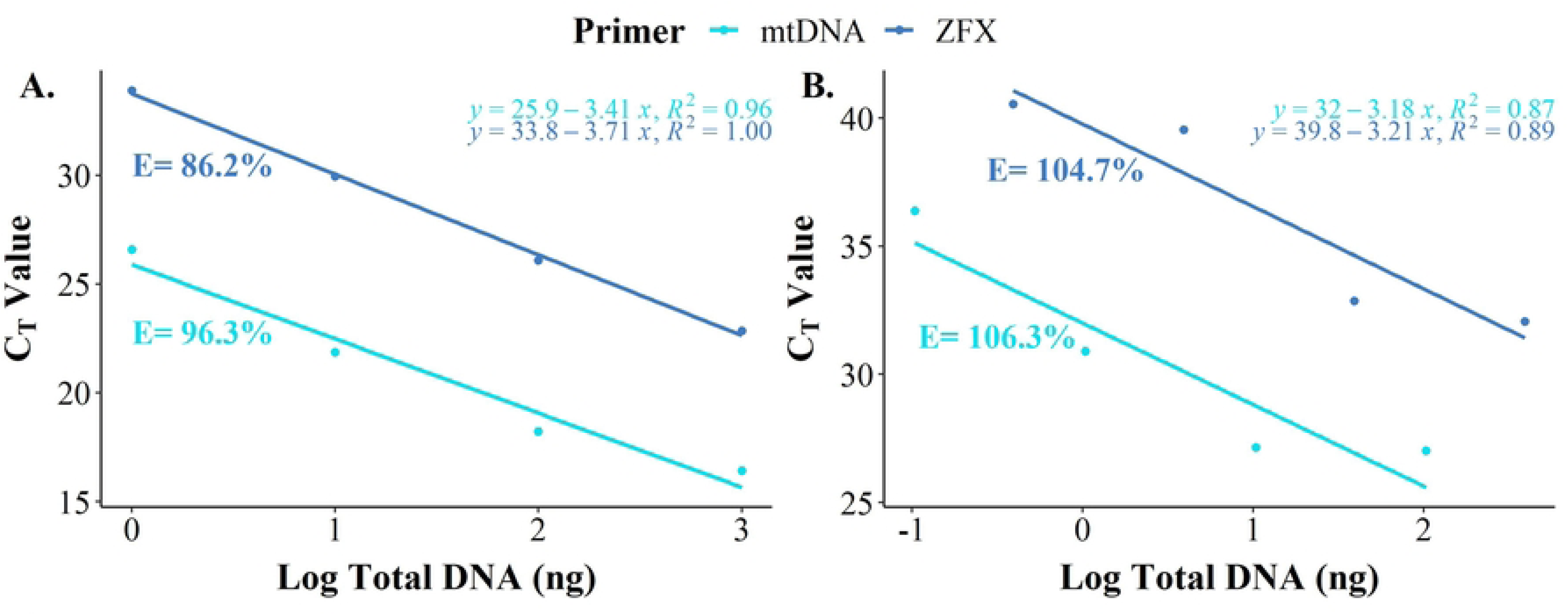
PCR dilution curves showing the efficiency (*E*) calculated in percentage for each primer sets. (A) The dilution curve of tissue DNA isolates extracted using the Qiagen DNeasy method. (B) The dilution curve of faecal DNA isolates extracted using the HV-CTAB-PCI method. The *E* showed slight inhibition for both primers, using faecal DNA isolates. The initial total DNA amount (ng) used for PCR reactions was determined from these dilutions.

### DNA extraction method comparison

The scrape-and-grind technique yielded significantly higher concentrations of total DNA (70.7 ± 5.3 ng/μL; mean ± standard error) compared to the scrape-only (34.9 ± 4.8 ng/μL, Tukey-HSD: T_14_ = 8.361, p < 0.001) and swabbing (20.4 ± 2.8 ng/μL, Tukey-HSD: T_14_ = 11.762, p < 0.001) techniques (Fig 4A). Total DNA concentrations, yielded using the scrape-only technique, were significantly higher than that of swabbing (Tukey-HSD: T_14_ = 3.401, p = 0.011). The scrape-and-grind technique also had higher inverse C_T_ values (0.040 ± 0.001) compared to scrape-only (0.033 ± 0.001, Tukey-HSD: T_14_ = 6.027, p < 0.001) and swabbing (0.035 ± 0.001, Tukey-HSD: T_14_ = 4.413, p = 0.002) techniques (Fig 4B). However, the inverse C_T_ values were not significantly different between scrape only and swabbing techniques (Tukey-HSD: T_14_ = 1.613, p = 0.273).

**Fig 4.**
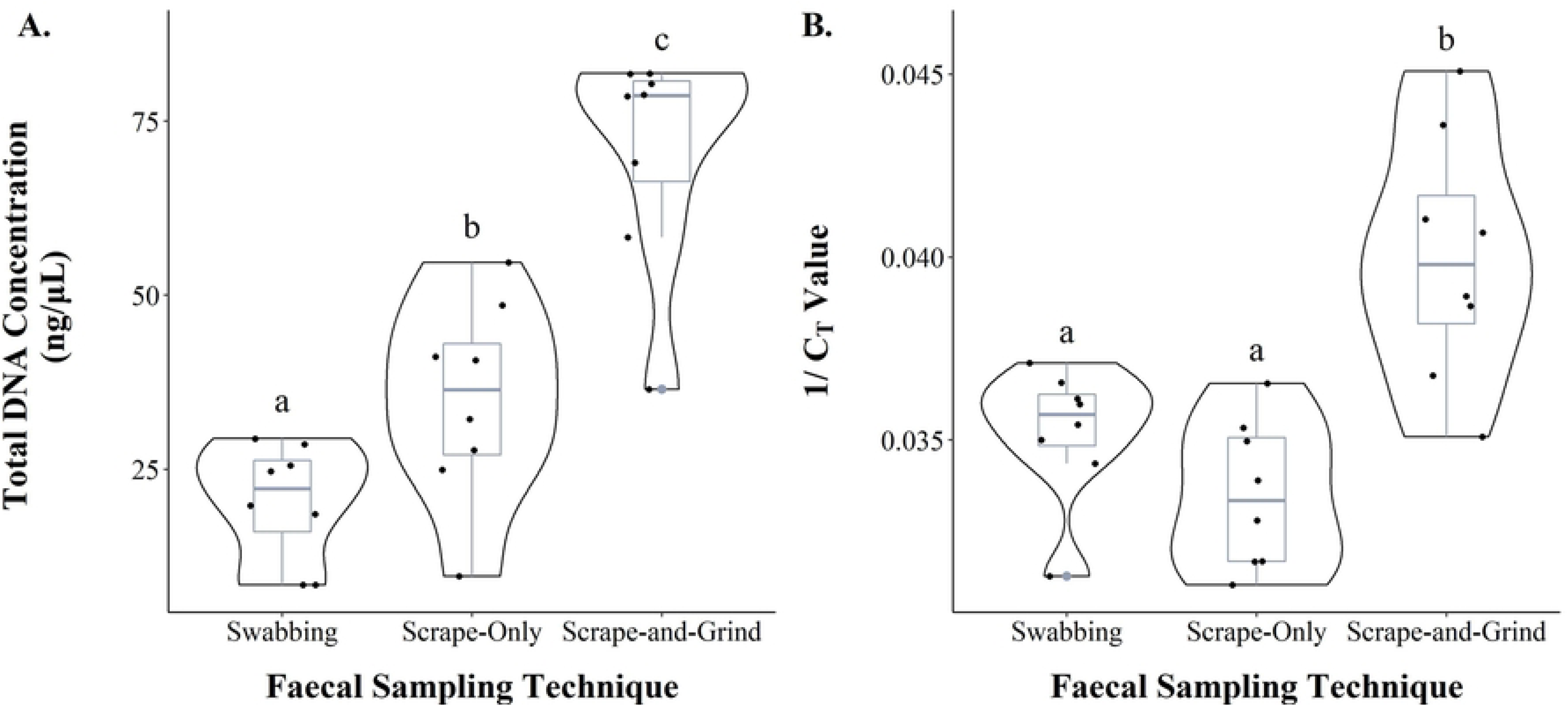
Comparison between faecal sampling and processing techniques to determine the best sampling protocol for DNA extraction. Violin plot shows the distribution of the data, whilst the box plot shows the first (Q1), second (Q2) and third (Q3) quartiles, minimum, maximum values, and outliers. All DNA was extracted using the QIAamp protocol. (A) Comparison of total DNA concentration extracted using each sampling and processing technique. (B) Comparison of the inverse C_T_ values from mtDNA amplification using faecal DNA extracted with each sampling and processing technique. Similar letters denote no statistical significance, while different letters denote statistical significance with an alpha value of 0.05.

Total DNA concentration extracted from faeces using HV-CTAB-PCI method (691.9 ± 51.9 ng/μL) was significantly higher than that extracted using QIAamp method (158.5 ± 15.8 ng/μL, Tukey-HSD: T_14_ = 11.701, p < 0.001, Fig 5A). The former method extracted more total DNA than control (145.4 ± 12.6 ng/μL, Tukey-HSD: T_14_ = 12.196, p < 0.001), but total DNA extracted using QIAamp method was not significantly different from control (Tukey-HSD: T_14_ = 0.496, p = 0.875). On average, the purity of faecal DNA extract was higher using QIAamp rather than HV-CTAB-PCI method (Table 5).

**Fig 5.**
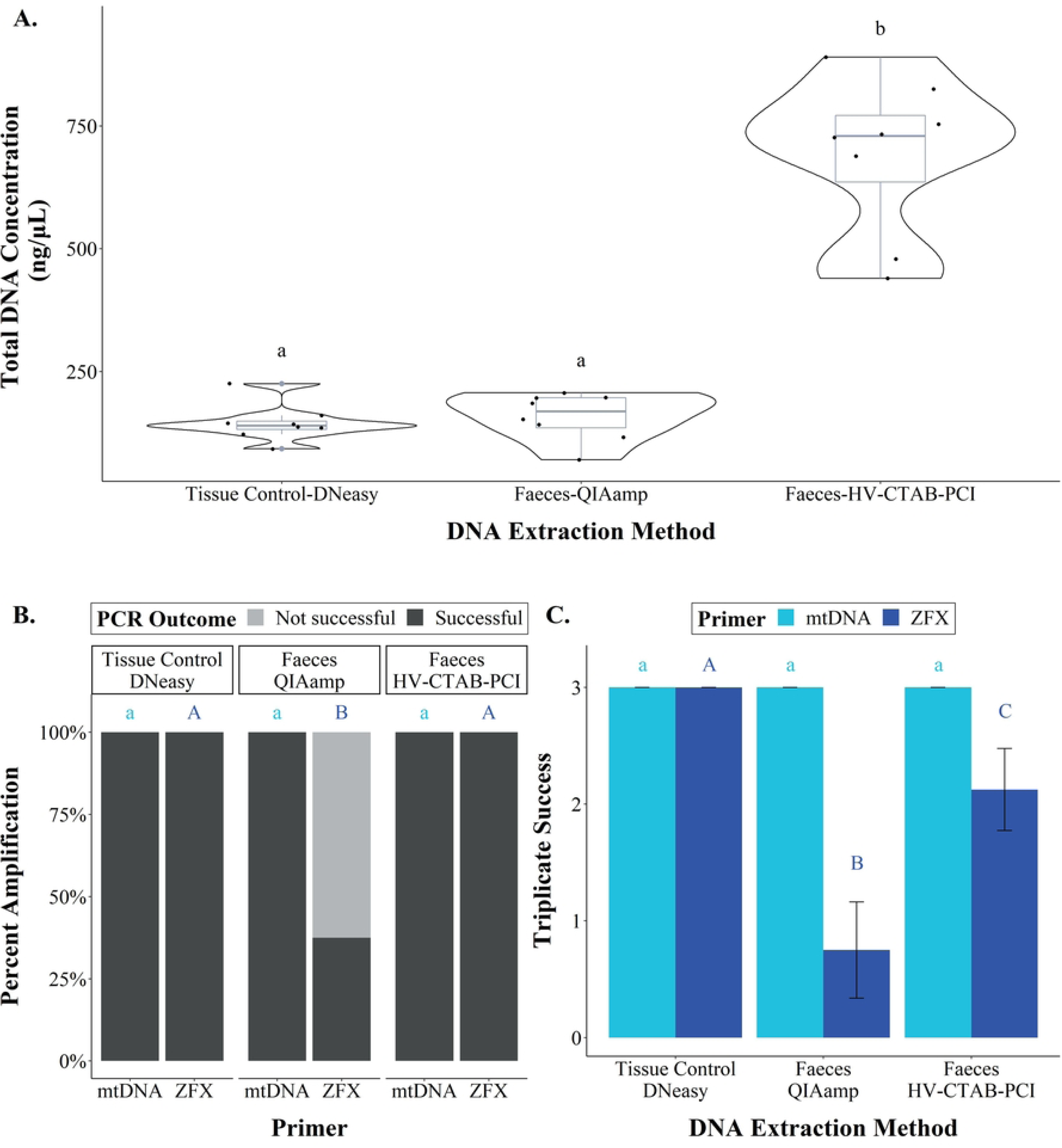
Comparisons between two DNA extraction methods and a tissue control. (A) A violin plot with box plot showing the total DNA concentration of each DNA extraction method. (B) A stacked bar graph showing the percent amplification that was successful (PCR success) and unsuccessful between all DNA extraction methods for each primer. (C) A bar graph showing the mean ± standard error of the number of replicates that amplified in triplicate for each DNA extraction method. Similar letters (with same colour and letter case) denote no statistical significance, while different letters denote statistical significance (α= 0.05). Different statistical comparisons made were indicated by different colour and letter case.

**Table 5.** Purity (A260/280 and A260/230) values of DNA extracted for each sample type using the Qiagen kits and a newly developed HV-CTAB-PCI method.

| DNA Extraction Method | Sample Type | A260/280 (mean ± std error) | A260/230 (mean ± std error) |
| --- | --- | --- | --- |
| Qiagen DNeasy | Tissue | 2.12 ± 0.01 | 1.82 ± 0.04 |
| QIAamp | Faeces | 1.95 ± 0.01 | 1.27 ± 0.03 |
| HV-CTAB-PCI | Tissue | 2.06 ± 0.01 | 2.14 ± 0.01 |
| HV-CTAB-PCI | Faeces | 1.67 ± 0.01 | 1.06 ± 0.03 |

PCR success and triplicate success for mtDNA amplification were similar between all DNA extraction methods, as all replicates of all samples amplified successfully (Fig 5B, C). For ZFX amplification, PCR success of HV-CTAB-PCI method was the same as control, and PCR success of those two were significantly higher than that of the QIAamp method (χ^2^= 7.273, p = 0.026). All samples amplified in control and HV-CTAB-PCI method, but only 37.5% of faecal samples amplified in the QIAamp method (Fig 5B). The control had significantly higher triplicate success for ZFX amplification (Mean, x̄ amplification= 3/3) compared to HV-CTAB-PCI (x̄ amplification= 2.1/3, *W* test: Z= 1.86, p = 0.032) and QIAamp (x̄ amplification= 0.8/3, *W* test: Z= 2.69, p = 0.004) methods (Fig 5C). However, triplicate success of HV-CTAB-PCI method was higher than that of QIAamp method (*W*-test: Z= 1.86, p = 0.032).

### Effects of EELs and faecal layers sampled

Effects of sampling from different faecal layers did not depend on faecal EELs as all interactive models had higher AIC values compared to additive models (Table 6). Same results were obtained for LR-Chi-Square test (Deviance= 0.022 – 0.832, p > 0.842).

**Table 6.** The AIC values obtained for interactive and additive models fitted in R for different variables compared using a dual-direction stepwise model selection.

| Response variable | Type of model | AIC values |
| --- | --- | --- |
| Total DNA concentration | Interactive | 349.98 |
|  | Additive | 346.74 |
| Log relative amount of mtDNA | Interactive | 930.97 |
|  | Additive | 929.56 |

#### Impacts of faecal layer sampled

For faeces of all EELs, total DNA concentration extracted from the outer surface of faeces was significantly higher than that extracted from their inner core (Tukey-HSD: T= 5.601, p < 0.001 for all faeces; Fig 6A). The outer surface of *ex*-EE-L1 faeces had at least 2.87× more mtDNA than their inner core, while the outer surface of *in*-EE-L3, L4, and L5 faeces had at least 0.14×, 0.65×, and 0.95× more mtDNA than their inner core, respectively (Fig 6B). On average, the outer surface of all stools had at least 0.16× more dugong mtDNA than the inner core of faeces. However, none of the differences were significant (Tukey-HSD: T= 1.222, p = 0.919 for all faeces). PCR success (Biased-Reduction-GLM: Z= 0.373, p = 0.709; Fig 6C) and triplicate success (Likelihood-ratio= 0.832, p = 0.842; Fig 6D) of mtDNA amplification were the same regardless of whether outer surface or inner core of faeces were sampled. As the layer of faeces sampled had no significant effect on the amplification success, only the outer surface of faeces was used for further comparisons.

**Fig 6.**
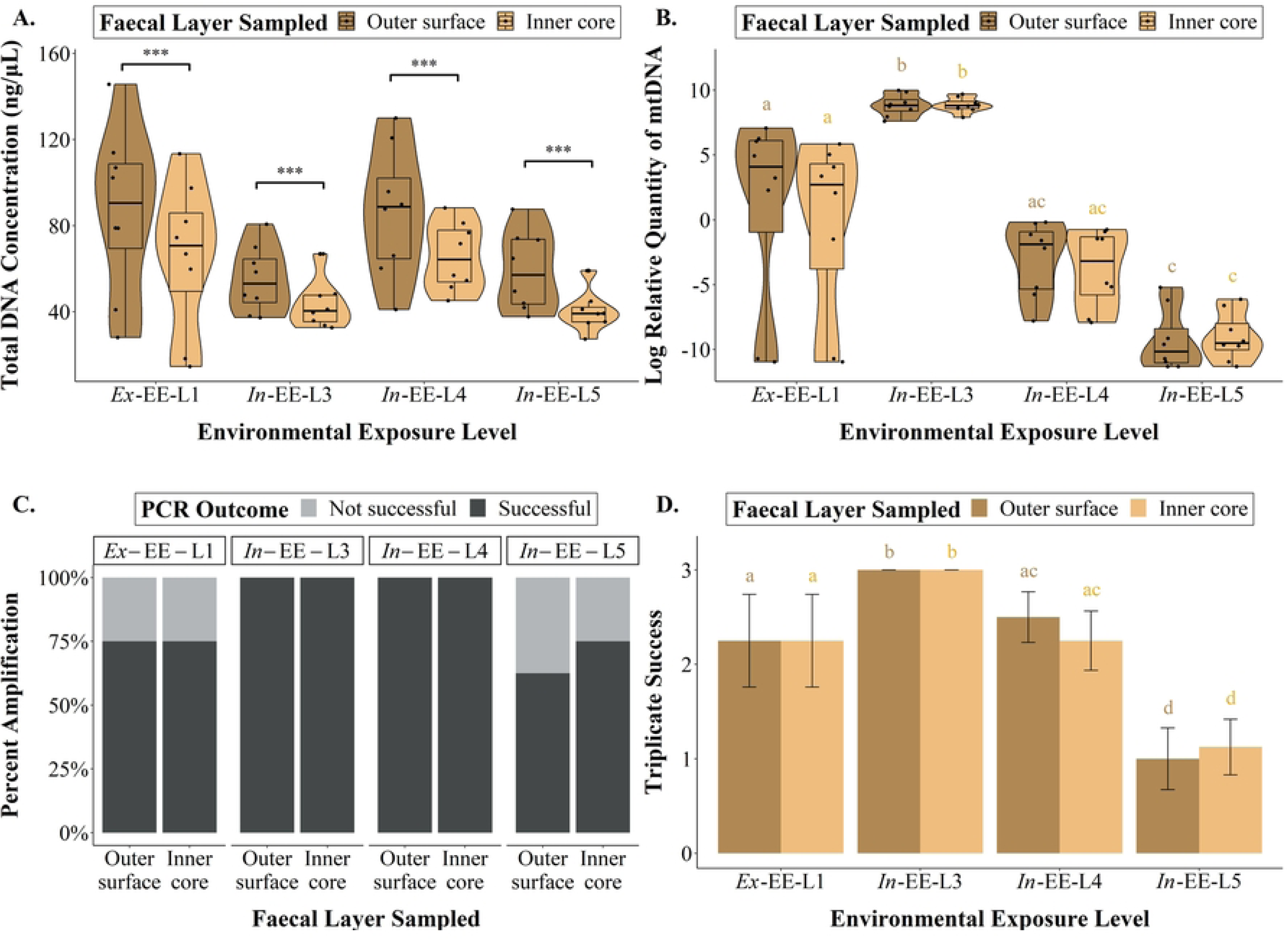
Comparison of the variables measured between outer surface and inner core of faeces across different environmental exposure levels (EELs) representing varying quality. (A) Total DNA concentration extracted from different layers of various faeces. No statistical significance was present between the faecal types, but the outer surface yielded a higher concentration of total DNA compared to the inner core for all faeces. (B) Log relative quantity of dugong mtDNA amplified from different layers of various faeces. (C) The percent mtDNA amplification that was successful and unsuccessful between different layers of various faeces. PCR success was statistically similar between all faecal types. (D) Triplicate success between different layers of various faeces. Similar letters denote non-significance, while different letters denote statistical significance (α= 0.05).

#### Impacts of environmental exposure

When the QIAamp method was used, the total DNA concentration extracted from *ex*-EE-L1 (87.0 ± 12.8 ng/μL), *in*-EE-L3 (55.2 ± 5.1 ng/μL), *in*-EE-L4 (86.5 ± 10.0 ng/μL), and *in*-EE-L5 (59.2 ± 6.06 ng/μL) faeces were not significantly different from each other regardless of the faecal layer sampled (Tukey-HSD: -2.877 <T_28_ < 0.970, p > 0.123; Fig 6A). The amount of dugong mtDNA extracted from *ex*-EE-L1 faeces was at least 46× less than that extracted from *in*-EE-L3 faeces (Tukey-HSD: T_30_ = -4.548, p < 0.001), but was 247,253× higher than that isolated from *in*-EE-L5 faeces (Tukey-HSD: T_30_ = 5.105, p < 0.001) (Fig 6B). Amount of dugong mtDNA was not significantly different between *ex*-EE-L1 and *in*-EE-L4 faeces (Tukey-HSD: T_29_ = 1.910, p= 0.246). However, amount of dugong mtDNA in *in*-EE-L3 faeces was at least 38,322× and 11,697,397× higher than that of the *in*-EE-L4 (Tukey-HSD: T_29_ = 6.604, p= < 0.001) and *in*-EE-L5 (Tukey-HSD: T_30_ = 14.405, p= < 0.001) faeces, respectively. The amount of dugong mtDNA was the same in *in*-EE-L4 and L5 faeces (Tukey-HSD: T_29_ = 3.049, p= 0.081).

PCR success for mtDNA amplification was similar between faeces of all EELs (Tukey-HSD: -1.583 < Z < 1.775, p > 0.285; Fig 6C). Despite insignificant results, all *in*-EE-L3 and L4 faeces amplified, but only 70% of *ex*-EE-L1 and 68.8% of *in*-EE-L5 faeces amplified successfully. However, *in*-EE-L3 faeces had higher triplicate success (x̄ amplification= 3/3) compared to *ex*-EE-L1 (x̄ amplification= 2.3/3, *W*-test: Z= 1.680, p= 0.046), *in*-EE-L4 (x̄ amplification= 2.4/3, *W*-test: Z= 2.530, p= 0.006) and *in*-EE-L5 faeces (x̄ amplification= 1.1/3, *W*-test: Z= 4.68, p < 0.001) (Fig 6D). Triplicate success of *ex*-EE-L1 faeces was not significantly different compared to that of *in*-EE-L4 faeces (*W*-test: Z= 0.444, p= 0.672) but was significantly higher than that of *in*-EE-L5 faeces (*W*-test: Z= 2.530, p= 0.006). The *in*-EE-L4 faeces also had a higher triplicate success than *in*-EE-L5 faeces (*W*-test: Z= 3.011, p= 0.001).

When the HV-CTAB-PCI method was used, the total DNA concentration extracted from *ex*-EE-L1 faeces (742.3 ± 70.3 ng/μL; mean ± standard error) was not significantly different to that extracted from *in*-EE-L2 faeces (559.4 ± 95.4 ng/μL, ANOVA: T_36_ = 1.934, p= 0.318), but was significantly higher than that extracted from *in*-EE-L3 (324.1 ± 19.8 ng/μL, ANOVA: T_36_ = 4.372, p= 0.001), *in*-EE-L4 (134.8 ± 14.4 ng/μL, ANOVA: T_36_ = 9.253, p < 0.001) and *in*-EE-L5 (198.3 ± 34.8 ng/μL, ANOVA: T_36_ = 7.597, p < 0.001) faeces (Fig 7A). Total DNA concentration of *in*-EE-L2 faeces was not significantly different compared to that of *in*-EE-L3 faeces (ANOVA: T_36_ = 2.369, p = 0.147), but was significantly higher that than of *in*-EE-L4 (ANOVA: T_36_ = 7.113, p < 0.001) and *in*-EE-L5 (ANOVA: T_36_ = 5.503, p < 0.001) faeces. The *in*-EE-L3 faeces also had higher concentration of total DNA compared to *in*-EE-L4 (ANOVA: T_36_ = 4.744, p < 0.001) and *in*-EE-L5 (ANOVA: T_36_ = 3.134, p = 0.027) faeces, but total DNA concentration between *in*-EE-L4 and L5 faeces was not significantly different (ANOVA: T_36_ = 1.610, p = 0.501).

**Fig 7.**
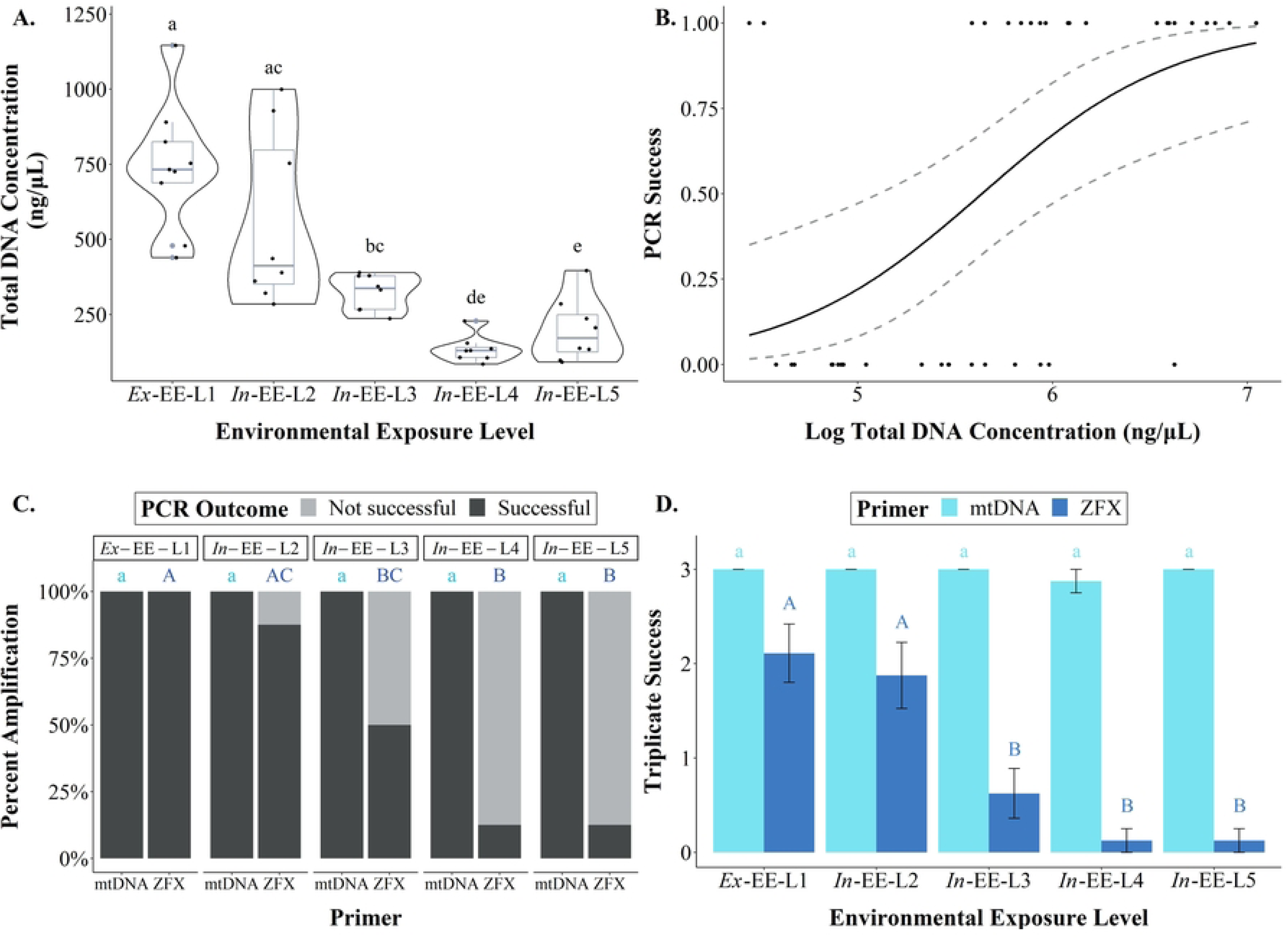
Comparison of variables measured in faeces with different environmental exposure levels (EELs) representing varying quality. (A) Total DNA concentration extracted from each faecal type. (B) Graph shows a positive correlation between PCR success and total DNA concentration. Lines represent mean ± standard error. (C) Percent amplification that was successful and unsuccessful for mtDNA and ZFX primers between different faecal types. (D) Triplicate success between the faecal types for both primers. Similar letters (with same colour and letter case) denote no statistical significance, while different letters denote statistical significance (α= 0.05). Different statistical comparisons made were indicated by different colour and letter case.

PCR success and triplicate success (Kruskal-Wallis: χ^2^= 4.125, p= 0.389) for mtDNA amplification were the same across faeces of all EELs, as all samples amplified successfully (Fig 7C, D). Mean triplicate success for faeces of all EELs was 3, except for *in*-EE-L4 faeces which was 2.9. For ZFX amplification, faeces with higher total DNA concentration tended to have higher PCR success (GLM: Z= 2.558, p=0.011; Fig 7B). PCR success of *ex*-EE-L1 faeces was similar to that of *in*-EE-L2 faeces (χ^2^= 1.195, p= 0.471), but was significantly higher than that of *in*-EE-L3 (χ^2^= 5.885, P= 0.032), *in*-EE-L4 (χ^2^= 13.39, p< 0.001), and *in*-EE-L5 (χ^2^= 11.46, p= 0.003) faeces (Fig 7C). All *ex*-EE-L1 faeces amplified, but only 87.5% and 50.0% of *in*-EE-L2 and L3 faeces amplified, respectively. Only 12.5% of both *in*-EE-L4 and L5 faeces amplified. PCR success of *in*-EE-L2 faeces was not significantly different compared to that of *in*-EE-L3 faeces (χ^2^= 2.618, p= 0.283), but was higher than that of *in*-EE-L4 (χ^2^= 9.00, P= 0.012) and *in*-EE-L5 faeces (χ^2^= 7.244, p= 0.015). PCR success of *in*-EE-L3, L4, and L5 faeces was not significantly different from one another (0.275 < χ^2^ < 2.618, p> 0.284). Triplicate success of *ex*-EE-L1 faeces (x̄ amplification= 2.1/3) was similar to that of *in*-EE-L2 faeces (x̄ amplification= 1.9/3, *W-t*est: Z= 0.711, p= 0.761), but higher than that of *in*-EE-L3 (x̄ amplification= 0.6/3, *W-*test: Z= 2.197, p= 0.014), *in*-EE-L4, and L5 (both x̄ amplification= 0.1/3, both *W-*tests: Z= 2.759, p= 0.003) faeces (Fig 7D). The *in*-EE-L2 faeces also had a higher triplicate success compared to *in*-EE-L3 (*W-*test: Z= 1.794, P= 0.036), *in*-EE-L4, and L5 (both *W-*tests: Z= 2.473, p= 0.007) faeces, but triplicate success of *in*-EE-L3, L4 and L5 faeces were not significantly different from each other (*W-*tests: Z < 1.029, p> 0.152).

### SNP amplifications in *in-ōceanum* faeces

All *in*-EE-L2 faecal samples amplified for all dugong SNP primers, except for primers Dug12 and Dug63, in which 50% and 75% of all samples amplified, respectively (Fig 8A). The SNP regions were successfully located and aligned, showing the form of allele possessed (S3 Fig). However, PCR success was not significantly different between amplification of different primer sets (χ^2^ < 5.33, p> 0.079; Fig 8A). Triplicate success of primer Dug12 (x̄ amplification= 1.3/3) was significantly lower than that of primers Dug50, Dug53, and Dug62 (All x̄ amplifications= 3/3, *W*-tests: Z= 1.694, p= 0.045) (Fig 8B). Triplicate success of primer Dug63 (x̄ amplification= 1.3/3) was equal to that of Dug12, but significantly lower than that of all other primers (All x̄ amplifications > 2.8/3, *W*-tests: Z > 2.115, p < 0.017).

**Fig 8.**
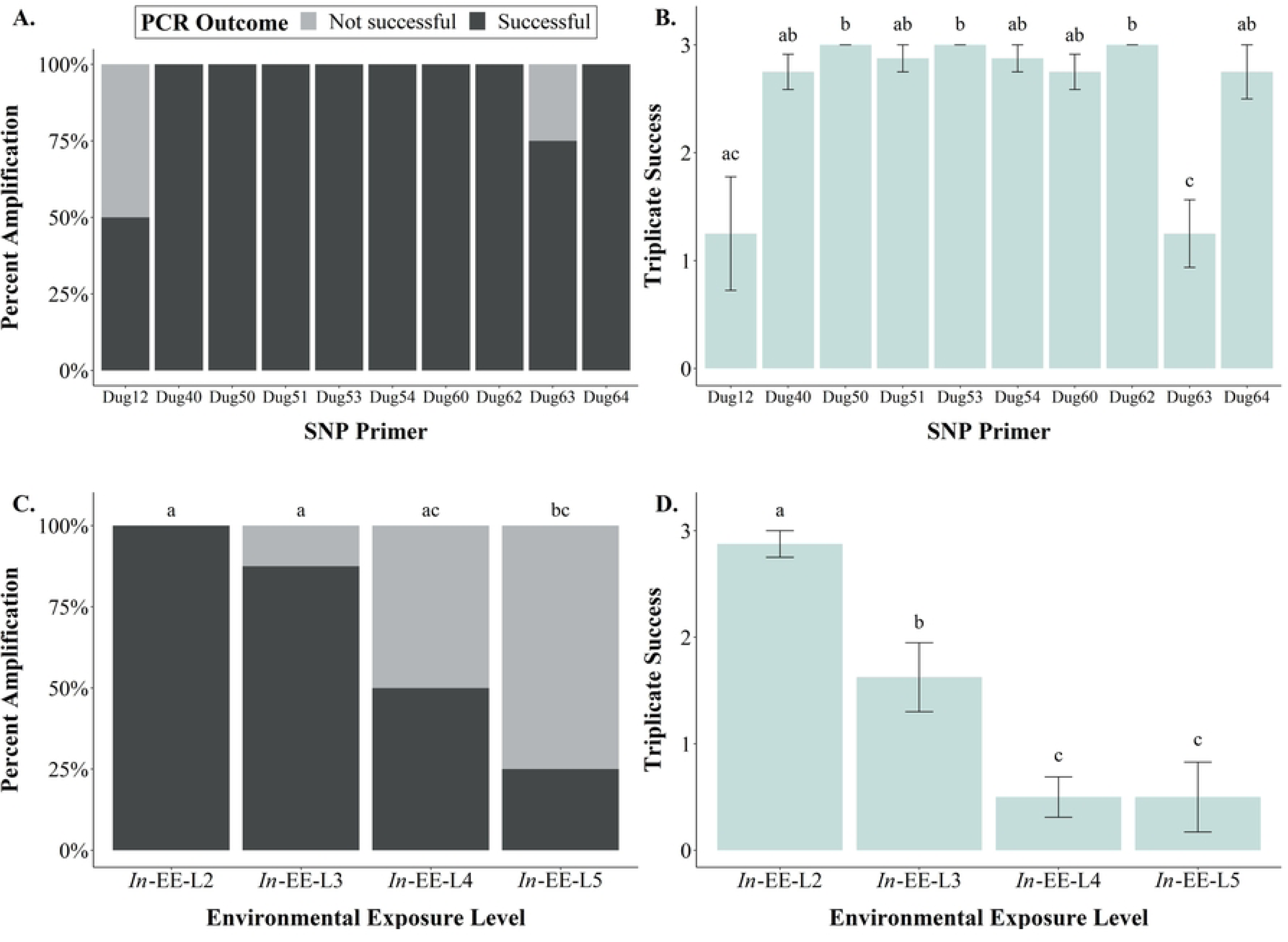
PCR success and triplicate success between SNP primers and faeces with different environmental exposure levels (EELs), representing its quality. (A) PCR success of different SNP primers amplified. No statistical significance exists between primers used. (B) Triplicate success of different SNP primers amplified. (C) PCR success between faeces of different EELs. (D) Triplicate success between faeces of different EELs. Similar letters denote no statistical significance, while different letters denote statistical significance (α= 0.05).

For amplification of SNP primer Dug54 in all *in-ōceanum* faeces, the PCR success of *in*-EE-L2 faeces was significantly higher than that of *in*-EE-L5 faeces (χ^2^= 9.600, p= 0.008), but similar to that of *in*-EE-L3 and L4 faeces (χ^2^< 5.333, p > 0.077) (Fig 8C). All *in*-EE-L2 faeces amplified, and 87.5%, 50.0%, and 25.0% of *in*-EE-L3, L4, and L5 faeces amplified, respectively. The *in*-EE-L3 faeces also had a higher PCR success compared to *in*-EE-L5 faeces (χ^2^= 6.349, p= 0.042), but it had a similar PCR success as *in*-EE-L4 faeces (χ^2^= 2.618, p= 0.283). The PCR success of *in*-EE-L4 and L5 faeces was not significantly different (χ^2^= 1.067, p= 0.603). The triplicate success of *in*-EE-L2 faeces (x̄ amplification= 2.9/3) was significantly higher than that of the *in*-EE-L3 (x̄ amplification= 1.6/3, *W-*test: Z= 2.382, p= 0.009), *in*-EE-L4, and L5 faeces (both x̄ amplification= 0.5/3, both *W-*tests: Z= 2.929, p=0.002) (Fig 8D). The *in*-EE-L3 faeces also had higher triplicate success compared to *in*-EE-L4 (*W-*test: Z= 1.937, p= 0.026) and *in*-EE-L5 faeces (*W-*test: Z= 1.706, p= 0.044). However, triplicate success of *in*-EE-L4 and L5 faeces was the same (*W-*test: Z= 0.438, p= 0.669).

## Discussion

This study demonstrated that mitochondrial and nuclear markers can be amplified from dugong faeces, using a novel ‘High volume-CTAB-PCI’ (HV-CTAB-PCI) DNA extraction method. Numerous modifications of the 2CTAB/PCI method by Vallet *et al* [18] resulted in this efficacious method for DNA isolation from a large quantity of faecal material. The amplification success of mtDNA was found to be similar regardless of whether faecal material was sampled from the outer surface or inner core of a stool. However, the period to which faeces were exposed to the environment had significant impacts on amplification success of nDNA markers. Although the amplification success of mtDNA was adequate with the QIAamp method, the use of HV-CTAB-PCI resulted in improvements, especially for eroded faeces (*in*-EE-L5). Regardless of the method used, nuclear markers amplified better in fresher faeces than eroded ones. Therefore, the hypothesis suggesting that QIAamp method would enable both mtDNA and nDNA amplifications when sampling from a stool’s outer surface was rejected; the hypothesis suggesting increased efficacy of amplification in fresher faeces was supported by these results.

### Development of HV-CTAB-PCI method

The HV-CTAB-PCI method developed in this study was cost-effective, user-friendly, and highly efficacious at isolating sufficient quantity and quality of dugong DNA from scats. Although mtDNA amplification was highly successful using DNA extracted following the QIAamp protocol, ZFX amplification was inconsistent, even for fresh faecal samples. This contrasts with Takoukam Kamla’s [32] results showing that the QIAamp protocol enabled amplifications of both microsatellites and mtDNA from manatee scats. In his study, pre-amplification enhanced the amount of manatee DNA that was originally extracted, thus improving the subsequent PCRs for nDNA amplification. However, pre-amplification in pilot experiments (including touchdown protocol) conducted in this study yielded no ZFX (nDNA) amplifications in the final PCRs. This may suggest that there is only a scant amount of DNA in dugong faeces. If the amount of dugong DNA originally extracted was of a negligible proportion compared to the robust quantities of exogenous DNA, then such a dilute presence of dugong DNA would render an infinitesimal probability that dugong DNA is pipetted into each well of the PCR plates [55]. If no dugong DNA was pipetted into the PCR reactions, then pre-amplification would not increase the quantity of target DNA, thus resulting in non-amplification of target DNA in subsequent PCRs. Another notable contributor to Takoukam Kamla’s success was likely the addition of a purification step post-DNA isolation, removing PCR inhibitors that may have been co-extracted with the DNA [32]. However, the consistent amplification of dugong mtDNA and results from dilution experiments performed in this study showed only low levels of PCR inhibition. Consequently, an additional purification step was not utilised in this study, in an attempt to minimise DNA loss [56]. Furthermore, this study showed a positive correlation between concentration of total DNA extracted and PCR success. This again hinted at a problem of scarcity of DNA in dugong faeces rather than issues of PCR inhibitors. As dugongs feed on a more restricted diet [35] compared to manatees [33], it is possible that there are fewer and less diverse plant secondary metabolites (e.g., tannins) to inhibit PCR in dugong faeces. Additionally, unlike the open ocean where dugongs are found, the estuarine and freshwater water bodies that manatees inhabit often contain high concentrations of tannins due to the decomposition of plant material from the surrounding forests and mangroves. Furthermore, the surface of dugong faeces is generally smoother and less fibrous than that of manatees. This may affect the relative quantity of epithelial cells found within dugong faeces, as the rougher, harder scats of manatees may stimulate more mucus secretion that entraps exfoliated cells [13].

A simple way to increase total DNA extracted from faeces is to increase the total amount of faecal material used for DNA extraction. This study utilised 220 mg of dugong faeces for DNA extraction with the QIAamp protocol as this is the maximum amount recommended by the manufacturer. Despite the retention of a higher volume of supernatant that contains DNA and the increase in subsequent volumes of extraction buffers used, this method did not isolate sufficient target DNA for consistent ZFX amplification in this study. Takoukam Kamla [32] managed to use a slightly larger amount (300 – 1260 mg) of manatee faeces without increasing the volumes of any reagents used. However, that might have contributed to the lower purity of those DNA extracts since there may have been insufficient reagents to efficiently clean and protect the extracted DNA, making an extra purification step essential. Since target DNA insufficiency was a problem in this study, the HV-CTAB-PCI method was developed to enable a more economical higher volume extraction of faeces. This method was modified from Vallet *et al*’s [18] 2CTAB/PCI protocol that was successful in extracting DNA from faeces of herbivorous primates. As the reagents used were self-prepared and the function of each chemical was known, this new method provided the flexibility to modify usage and concentrations of any reagents to suit DNA extraction from dugong faeces.

This novel HV-CTAB-PCI method is markedly different to the 2CTAB/PCI approach. The minimum effective sample-to-reagent ratio used in this method (1:5) was first determined through a 1 mL-by-1 mL addition of Lysis Buffer 1 to 1 g of faecal sample. This differs from the ratios used in the 2CTAB/PCI approach [18], which would require large volumes of reagents, when scaled up, to extract DNA from 1 g of faeces. Instead of an overnight incubation, cell lysis duration was reduced to 3 h to shorten the DNA extraction procedure. To separate DNA from proteins and other impurities, an effective equal PCI-to-supernatant ratio (1:1) was utilised [57]. Proteins were digested using protease instead of the widely used Proteinase K in the HV-CTAB-PCI method as protease was found to be effective at removing proteins, and costs less than the latter enzyme. As the presence of RNA did not present problems in PCR amplifications, RNAse was not used in this newly developed protocol. To minimise DNA loss, the DNA pellet retrieved at the end of extraction was washed once only [56]. Despite a reduced washing frequency, PCR was not significantly inhibited and the amplification successes of mtDNA and ZFX were higher using this new protocol compared to the QIAamp method. This again showed that PCR inhibitors were unlikely to be a major issue within dugong faeces.

### Sampling from different layers of faeces of varying quality

Amplification success and relative quantity of mtDNA did not differ between DNA extracted from the outer surface and inner core of dugong stools. This result was surprising as faecal DNA extractions in most molecular scatology studies have sampled the outer surface of a scat only, e.g., [8, 14]. This study concurs with Stenglein *et al* [58] which found no difference in PCR success and error rates of DNA extracted from outer and inner layers of brown bear scats. Although lower allelic dropout and higher genotyping success rates were achieved when DNA from the outer surface of stools was used [58], it would be improper to conclude that most of the target animal cells are located on the outer surface as those measures only reflect the quality of DNA extracted. As the total DNA yield tended to be higher when outer surface was sampled, it is possible that the small sample size (total n= 32) in this study limited the detection of significance between outer and inner layers of faecal stools. However, it is likely that most of the DNA extracted was from organisms that were not of interest (e.g., bacteria, plants) since the relative amount of dugong mtDNA was the same regardless of layers sampled. Instead, we hypothesise that the target animal cells and therefore their DNA would be heterogeneously distributed throughout the faecal mass. Since faeces are unformed in the upper regions of the colon, exfoliated epithelial cells from those regions would likely be incorporated within the faecal mass through peristaltic contractions of the colon [12, 59].

The degree to which voided dugong faeces were exposed to the environment was found to significantly influence amplification success and relative amount of recovered DNA. Comparing mtDNA amplifications, highly eroded stools were found to amplify poorly compared to fresher ones. Eroded faeces were anticipated to amplify poorly as DNA degrades over time [60]. Epithelial cells undergo apoptosis as they detach from the basement membrane [61], where enzymes that degrade DNA break down the phosphodiester bonds that form the backbone of DNA [62]. This breaks the DNA into progressively shorter fragments and eventually into nucleotides [62]. The new HV-CTAB-PCI method, however, greatly improved mtDNA amplification, allowing consistent amplification even in eroded faecal samples. As faecal DNA is often highly degraded, the extraction of DNA from a larger amount of faecal material, used in the HV-CTAB-PCI method, can increase the chance for more target DNA of sufficient length or quality to be extracted, thus allowing for a higher amplification success. As such, this newly developed DNA extraction method also enabled the comparison of ZFX amplifications in faeces of different environmental exposure levels (EELs). The amplification of ZFX marker was found to be highest in freshest faeces (*ex*-EE-L1) compared to older ones. Interestingly, the amplification success and relative quantity of mtDNA were found to be higher in fresh *in*-*ōceanum* faeces (*in*-EE-L3) compared to the *ex-ōceanum* faeces (*ex*-EE-L1), in contrast to what one might expect. The amount of amplifiable target DNA is influenced by the physical loss and chemical degradation (e.g., fragmentation) of DNA. Although the loss of target DNA could be higher when faeces is transported in the ocean, sea salts such as Ca^2+^ and Mg^2+^,which DNA has a higher affinity for compared to Na^+^ [63, 64], may help to stabilise or preserve the integrity of DNA [64,65,66]. Therefore, the higher relative quantity of mtDNA in *in*-EE-L3 faeces may be due to preservative effects of salt. However, the higher DNA loss in *in*-EE-L4 and L5 faeces may have outweighed any preservative effects on DNA, leading to the observed results. Although it remains unknown why the ZFX amplifications failed to reproduce the same trend, it is possible that the structural differences (circular versus linear) or the cellular location of mtDNA versus nDNA may have influenced their degradative vulnerability [67].

Although this study achieved a low amplification success in more eroded faeces, the chance of success may be enhanced by scaling up the amount of starting faecal material used for DNA extraction. If that fails, separation of host DNA from bacterial DNA could be trialled using the host-DNA enrichment method by Chiou and Bergey [68] as this may improve the purity of DNA extracted. However, whole genome amplification may need to be performed prior to enrichment if the amount of target DNA extracted is insufficient for an efficient separation. Alternatively, host epithelial cells could be separated from a large proportion or the entire faecal mass using the methodology developed by Matsushita et al. [69] and DNA extraction could then be directly performed on those cells.

### SNP amplifications using faecal DNA

This study showed a proof-of-concept for the potential utilisation of faecal DNA in population genetic studies of dugongs, as all ten SNPs were successfully amplified using the dugong faecal DNA extracted with the new HV-CTAB-PCI method. The SNP primers did not significantly influence amplification success, but interestingly, primers Dug12 and Dug63 amplified less than other SNP primers. One plausible explanation is that those target sites may be more sensitive to DNA degradation. For example, Johnston *et al* [70] showed that regions where SNP markers showed polyploidy due to historical genome duplication in Atlantic salmon had higher sensitivity to DNA degradation and thus had lower genotyping success. Alternatively, the primers used in this study that were designed for those target sites may not have been optimal. The impact of DNA degradation was also reflected in SNP amplifications as fresh faeces amplified more successfully than eroded ones. Since SNP amplicons are generally shorter than microsatellites [30], the difficulty for SNP amplification in eroded faeces may indicate that the DNA extracted from these faeces was too degraded or fragmented for primers to bind and amplify at the target site. That said, only one SNP marker (Dug54) was used to compare the amplification success between faeces of different EELs which limits the extent of inferences that can be made.

## Conclusion

This research has successfully pioneered a novel DNA extraction protocol (HV-CTAB-PCI) for an herbivorous marine mammal, the dugong, and has demonstrated the most efficient sampling methodologies to maximise retrieval of target DNA from faecal samples, to enable amplification of both mitochondrial and nuclear markers used in genetic population studies. This study also provided a preliminary idea on the distribution of sloughed epithelial cells and thus target DNA within faecal stools, which may help researchers increase the efficiency and success for target DNA recovery. The main limitation of this research was the inability to isolate DNA of sufficient quality from the oldest, most eroded stools for amplification of target nDNA. Despite that, multiple suggestions have been made to increase the chances for DNA retrieval. As the first study to demonstrate consistent amplification of nDNA from dugong faeces, the development of the HV-CTAB-PCI method will facilitate non-invasive genetic studies in areas where direct sampling is unfeasible. For instance, faecal samples have been collected from some remote and turbid water regions in northern Queensland where dugongs could not be sampled directly. These samples may now be analysed to gain a fuller understanding on the population structure of dugongs along the entire eastern Queensland coast. Future application of this approach could include broad-scale population genetic studies on dugongs throughout the Indo-Pacific region where resources for direct sampling are typically unavailable but obtaining samples for genetic analysis is critical.

## Acknowledgements

Thanks to the UQ Dugong Team for assistance in sample collection, especially Helen Peereboom. Morgan McCarthy provided SNP amplicon sequences, Jennifer Seddon advised on SNP analysis, Simone Blomberg advised on statistics, while Seth and Matthew Kazmer assisted with hand-drawn illustrations. Any use of trade, firm, or product names is for descriptive purposes only and does not imply endorsement by the U.S. Government.

## Supporting information

**S1 Fig. Flowcharts summarising the experimental designs and approaches taken in this study.**

(A) The first stage of this study targeted the development of a DNA extraction protocol for dugong faeces. (B) The second stage of this study assessed the influence of sampling between outer surface and inner core of faeces, in addition to impacts of faecal environmental exposure, on the amplification success of mtDNA and ZFX markers. (C) The final stage of this study sought to amplify ten SNP markers to show a proof-of-concept that population genetic markers used for genetic studies of dugongs can be amplified using their faecal DNA. The impacts on amplification were compared between different *In-ōceanum* faeces.

**S2 Fig. Results from a pilot study conducted to determine initial amount of total DNA (10 and 100 ng) to be added into qPCR assay for comparison of mtDNA amplification between outer surface and inner core of faeces with different EELs.**

(A) Stacked bar charts of PCR success for the different initial total DNA used. No difference was found between the initial DNA amount used (χ^2^= 0.381, p= 1.000). (B) Violin plots incorporating box plots of triplicate success for the different initial total DNA. No difference was found between the total DNA used (Kruskal-Wallis χ^2^= 0.804, df= 1, p= 0.370).

**S3 Fig. Alignment of SNP sequences amplified using ten dugong SNP primers with two forms of alleles known.**

Three random samples were chosen to be shown in the plot for each primer set. (A) Primer Dug12. (B) Primer Dug40. (C) Primer Dug50. (D) Primer Dug51. (E) Primer Dug53. (F) Primer Dug54. (G) Primer Dug60. (H) Primer Dug62. (I) Primer Dug63. (J) Primer Dug64.

